# Limits to Genomic Divergence Under Sexually Antagonistic Selection

**DOI:** 10.1101/591610

**Authors:** Katja R. Kasimatis, Peter L. Ralph, Patrick C. Phillips

**Affiliations:** Institute of Ecology and Evolution, 5289 University of Oregon, University of Oregon, Eugene, OR 97403

**Author notes:** **Corresponding Author:** Katja Kasimatis, Institute of Ecology and Evolution, 5289 University of Oregon, Eugene, OR 97403.

**Keywords:** antagonistic selection, sexual conflict, male-female divergence, intersexual *F*_*ST*_, genetic load

## Abstract

Since the autosomal genome is shared between the sexes, sex-specific fitness optima present an evolutionary challenge. While sexually antagonistic selection might favor different alleles within females and males, segregation randomly reassorts alleles at autosomal loci between sexes each generation. This process of homogenization during transmission thus prevents between-sex allelic divergence generated by sexually antagonistic selection from accumulating across multiple generations. However, recent empirical studies have reported high male-female *F*_*ST*_ statistics. Here, we use a population genetic model to evaluate whether these observations could plausibly be produced by sexually antagonistic selection. To do this, we use both a single-locus model with nonrandom mate choice, and individual-based simulations to study the relationship between strength of selection, degree of between-sex divergence, and the associated genetic load. We show that selection must be exceptionally strong to create measurable divergence between the sexes and that the decrease in population fitness due to this process is correspondingly high. Individual-based simulations with selection genome-wide recapitulate these patterns and indicate that small sample sizes and sampling variance can easily generate substantial male-female divergence. We therefore conclude that caution should be taken when interpreting autosomal allelic differentiation between the sexes.

## Introduction

Females and males use largely the same genome to produce distinct phenotypes and behaviors. This ubiquitous phenomenon requires an association between dimorphic phenotypes and their sexual environment (Kasimatis *et al.* 2017; Mank 2017b). Genes residing on a sex chromosome have a physical link to sex determination. Particularly, on heteromorphic sex chromosomes, the lack of recombination allows for selection to act in a sex-specific manner to optimize beneficial genes within each sex (Rice 1984, 1987; Charlesworth and Charlesworth 1980). Conversely, the shared genetic basis of autosomal genes constrains such sex-specific optimization of fitness. When autosomal-based traits have different optimal fitness values in each sex, then selection acts in a sexually antagonistic manner to push females and males in opposing directions in phenotype space (Rice and Holland 1997; Bonduriansky and Chenoweth 2009). However, recombination and meiotic segregation uncouple beneficial alleles from their sexual environment every generation, constraining the resolution of antagonism via the creation of separate female and male genomic pools. This homogenization process tethers together the evolutionary responses of the sexes and creates an inherent intersexual genomic conflict (reviewed in Bonduriansky and Chenoweth 2009; Kasimatis *et al.* 2017).

Identifying sexually antagonistic loci – particularly using reverse genomics approaches – has proved challenging. Initial studies calculated differentiation between females and males using Wright’s fixation index (*F*_*ST*_), and interpreted high values as evidence of sexually antagonistic selection. Empirical data from multiple taxonomic groups (Lucotte *et al.* 2016; Flanagan and Jones 2017; Wright *et al.* 2018; *Dutoit et al.* 2018) suggest that hundreds to thousands of SNPs have elevated male-female autosomal differentiation with outliers exceeding *F*_*ST*_ = 0.01 (Lucotte *et al.* 2016; Wright *et al.* 2018) and even approaching *F*_*ST*_ = 0.2 (Flanagan and Jones 2017). (Cheng and Kirkpatrick 2016) interpreted correlations of male–female *F*_*ST*_ with bias in gene expression to mean that many genes actively affect sex-specific viability in both humans and *Drosophila melanogaster*. Taken at face value, both the number of sexually antagonistic alleles and the degree of differentiation are striking. However, these results are difficult to evaluate as they suggest that there must be quite a large amount of selective mortality within each sex to create such high divergence within a generation (as noted by Cheng and Kirkpatrick 2016).

Several different processes could in principle generate divergence (or apparent divergence) between the sexes. First, sex biases in chromosome segregation through associations with the sex determining region could distort allele frequencies between the sexes. Over time, this segregation distortion can contribute to the generation of neo-sex chromosomes (Jaenike 2003; Kozielska *et al.* 2010) – particularly heteromorphic sex chromosomes – leading to sex-specific differentiation in the trivial sense that the locus is completely absent in one sex. Second, gametic selection resulting in a sex-specific fertilization bias could also distort allele frequencies (Joseph and Kirkpatrick 2004). Both of these processes occur during the gametic phase of the lifecycle and have long been recognized for their potential ability to distort segregation ratios within the sexes (reviewed in Immler and Otto 2018). In contrast, sexually antagonistic viability selection that occurs post-fertilization is a fundamentally different mechanism because there is no direct co-segregation of sex with the alleles under selection. Previous work on sexually antagonistic viability selection has largely focused on its potential role in maintaining genetic variation due to the sex-specific pleiotropic effects of the locus. In particular, Kidwell *et al.* (1977) laid out a framework for analyzing sexual antagonism that has widely been used in the field (Arnqvist 2011; Connallon *et al.* 2010; Connallon and Clark 2011; Patten and Haig 2009; Fry 2010). A little appreciated feature of the Kidwell model is that it tracks allele frequencies (rather than diploid genotype frequencies) in adults from each generation to the next. Although the model incorporates diploid selection, this sampling paradigm is sufficient because the “random union of gametes” model of mating only requires allele frequencies to generate diploid genotype frequencies in the next generation. However, this model simplification prevents the inclusion of other models of mating, such as assortative mating among genotypes.

In this paper, we will first build a model of sexually antagonistic viability selection, segregation, and transmission, extending the model of Kidwell *et al.* (1977) to include assortative mating. We use this model to evaluate how much between-sex differentiation is produced across a range of selection, dominance, and assortative mating parameters. Second, we use these results to evaluate the claims that the observed between-sex allelic differentiation is caused by sexually antagonistic viability selection. We then use simulation to test the conclusions of our deterministic model, as well as the role of sampling variance in generating loci with high between-sex differentiation. Both our single locus model and individual-based simulations with antagonistic loci distributed genome-wide indicate that antagonistic selection must be remarkably strong to produce non-negligible divergence between the sexes. Instead, simulations indicate that sampling variance is much more likely to account for extreme between-sex divergence and must therefore be explicitly included in any analyses of putative signatures of male-female divergence.

## Methods

### Model

Consider an autosomal locus in which are found two alleles: one female-beneficial (*A*_1_) and one male-beneficial (*A*_2_). Sexual antagonism results in a fitness cost to individuals carrying the allele favored in the other sex (Kidwell *et al.* 1977; Bodmer 1965). The life cycle is shown in figure 1. Each generation begins with zygotic frequencies equal in each sex, but then genotype-dependent survival results in different genotype frequencies in each sex at time of mating. The relative fitnesses of genotypes *A*_1_*A*_1_, *A*_1_*A*_2_, and *A*_2_*A*_2_ in females are 1:: 1 *− h_f_ s_f_*:: 1 *− s*_*f*_, where *s*_*f*_ is the cost of a female having the male-favorable allele and *h*_*f*_ is the dominance coefficient in females. (This is a model of viability selection, so here and below “fitness” refers to viability.) Writing the frequencies of the three genotypes in zygotes as *p*_11_(*t*), *p*_12_(*t*), and *p*_22_(*t*) at the start of generation *t*, the genotype frequencies in females after selection will then be proportional to *p*_11_(*t*), *p*_12_(*t*)(1 − *h*_*f*_ *s*_*f*_), and *p*_22_(*t*)(1 − *s*_*f*_), respectively. Similarly, the relative fitnesses of the genotypes *A*_1_*A*_1_, *A*_1_*A*_2_, and *A*_2_*A*_2_ in males are 1 *− s*_*m*_:: 1 − *h_m_s*_*m*_:: 1, and the genotype frequencies in males after selection are proportional to *p*_11_(*t*)(1 − *s*_*m*_), *p*_12_(*t*)(1 − *h_m_s*_*m*_), and *p*_22_(*t*), respectively.

**Figure 1:**
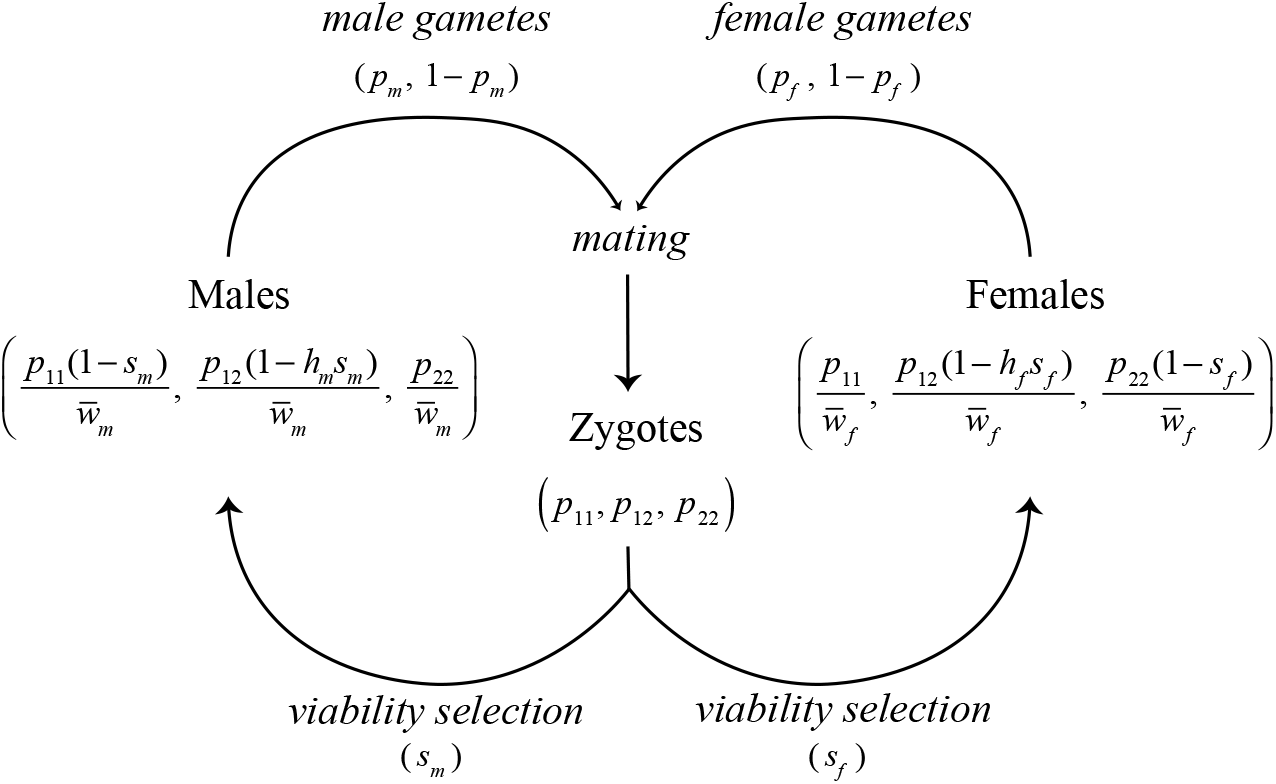
Lifecycle of the model. Zygotes are subject to sexually antagonistic viability selection (*s*_*m*_ and *s*_*f*_), perturbing allele frequencies in adults in a sex-specific manner. Sex-specific adult allele frequencies are given in Equations 1 and 2, where 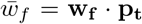 and 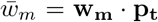. Surviving adults produce gametes of each allele type in frequencies corresponding to Equations 1 and 2. At this time meiotic segregation breaks the association between the locus and sex. Females and males mate with frequencies proportional to the mate choice matrix (**M**) to produce the zygote pool in the next generation. Kidwell *et al.* (1977) gives the recursion for the allele frequencies in gametes (*p*_*m*_, *p*_*f*_), under the assumption of random, genotypeindependent mating.

Therefore, the frequency of the female-beneficial allele in females post-selection, which we denote *p*_*f*_ (*t*), is

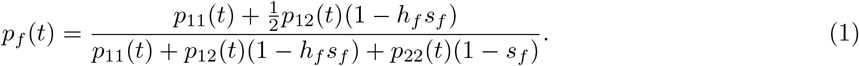

The same quantity for males is:

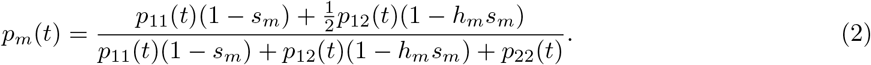

In a deterministic model of non-overlapping generations without gamete-specific selection, the genotype frequencies in the next generation are determined by the frequency of gametes joining from each of the nine possible mating combinations weighted according to mate choice. We parameterize mate choice using a matrix whose rows are indexed by male genotypes and columns by female genotypes, such that *M*_*ij*_ is the frequency of pairings of male genotype *i* with female genotype *j* relative to that expected under random mating. We focus on three common mating scenarios by structuring the mate choice matrix as:

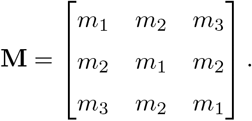

Under random mating, each pairing occurs with equal likelihood (*m*_1_ = *m*_2_ = *m*_3_). Positive assortative mating by genotype occurs when females and males with the same genotype mate more frequently than those with different genotypes (*m*_1_ *> m*_2_ = *m*_3_). Conversely, disassortative mating by genotype – or positive assortative mating by fitness – occurs when *A*_1_*A*_1_ individuals mate with *A*_2_*A*_2_ individuals (*m*_1_ = *m*_2_*< m*_3_). The genotype frequencies in the next generation can be concisely calculated with some matrix algebra. Let **w_f_** = (1, 1−*h*_*f*_ *s*_*f*_, 1*−s*_*m*_) and **w_m_** = (1−*s*_*m*_, 1−*h*_*m*_*s*_*m*_, 1) be the vectors of relative fitnesses in females and males respectively. Then, define the 3 *×* 3 matrix of fitness-weighted mate pairings, **F**, so that for each pair of genotypes *a* and *b*, the entry **F**_*ab*_ = *w*_*m*_(*a*)*M_ab_w*_*f*_ (*b*). In other words, **F** = diag(*w*_*m*_)*M* diag(*w*_*f*_), where diag(*w*_*m*_) denotes the matrix with **w_m_** on the diagonal and zeros elsewhere. Finally, define *β* = diag(1, 1*/*2, 0) and *γ* = diag(0, 1*/*2, 1). Then, the vector of frequencies of each genotype among zygotes (before selection) in the next generation can be calculated using the current frequencies as a weighted sum over possible mating pairs:

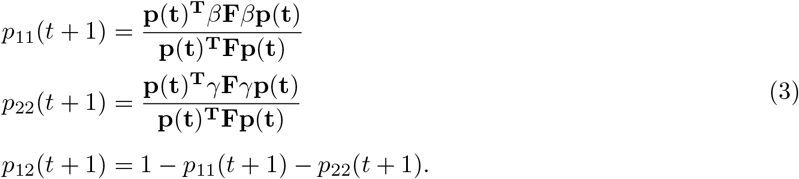

Here, **p**(**t**) = (*p*_11_(*t*)*, p*_12_(*t*)*, p*_22_(*t*)) is the column vector of genotype frequencies and **p**(**t**)^*T*^ is its transpose. This set of equations can be derived by noting that the relative frequencies of *A*_1_*A*_1_, *A*_1_*A*_2_, and *A*_2_*A*_2_ genotypes produced in the next generation are **p**(**t**)^*T*^ β**F***β***p**(**t**), **p**(**t**)^*T*^ (*β***F***γ* + *γ***F***β*)**p**(**t**), and **p**(**t**)^*T*^ γ**F***γ***p**(**t**), respectively; since *β* + *γ* = *I*, the identity matrix, these sum to **p**(**t**)^**T**^**Fp**(**t**), the denominator in equations (3).

We used Mathematica v11.1.1.0 (Wolfram Research, Inc.) to find the equilibria of this system and determine stability of those equilibria. The complete notebook is provided in File S1.

### Within-generation statistics

Sex-specific viability selection creates differences in allele frequencies between the sexes each generation. We can therefore quantify the effects of sexually antagonistic selection using the male-female *F*_*ST*_ statistic, which we calculate as the squared difference in allele frequencies between sexes, normalized by the total heterozygosity across sexes (Cheng and Kirkpatrick 2016; Wright 1951):

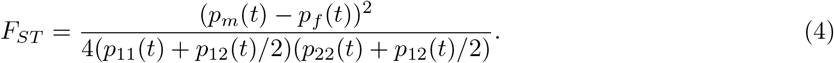

Sex-specific selection creates divergence between the sexes by increasing the frequency of the beneficial allele in each sex. Therefore, at the population level, this opposing action of section skews genotype frequencies away from Hardy-Weinberg equilibrium. The deviation from Hardy-Weinberg equilibrium within the population due to sex-specific effects can be quantified using Wright’s *F*_*IS*_ statistic (Wright 1951):

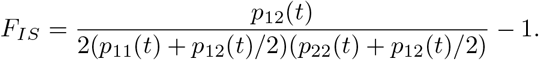

A population fitness cost due to sexual antagonism (i.e., genetic load), is generated each generation. Within each sex, the genetic load is the difference between the maximum possible fitness and the mean fitness (Haldane 1937). The population’s average genetic load (*L*) is the average of the loads for each sex (assuming an equal sex ratio), which is given by:

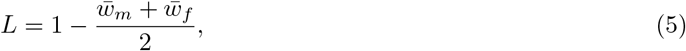

where 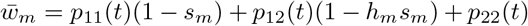 and 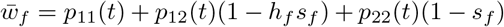.

### Simulations

We used R (R Core Team 2018) (File S2) to simulate allele frequency dynamics at a single locus in a population subject to selection and drift. During viability selection each generation, each individual survived with probability equal to their (sex- and genotype-dependant) fitness. Then, the genotype frequencies within each sex were multiplied to give the matrix of relative frequencies of possible mating pairs, which was further weighted by the mate choice matrix. To generate the next generation, a fixed number of mating pairs are sampled from this distribution, and offspring are produced by random choice of parental alleles.

We also implemented simulations with sexually antagonistic selection acting at many loci, genome-wide with SLiM v3.1, an evolution simulation framework (Haller and Messer 2019) (recipes in File S3). Individuals each had a genome of 100 Mb, a uniform recombination rate of 10^*−*8^ per nucleotide, and a mutation rate of 10^*−*10^. All mutations are sexually antagonistic (we do not simulate neutral variation): each new mutation was beneficial in a randomly chosen sex and detrimental in the other, with selection coefficients drawn independently for each sex from a Gaussian distribution with mean zero and standard deviation 0.01. Each mutation also had dominance coefficients drawn independently for each sex from a uniform distribution between 0 and 1. The model had overlapping generations: each time step, first viability selection occurred (with probability of survival equal to fitness), followed by reproduction by random mating. The number of new offspring was chosen so that the population size fluctuated around 10,000 diploids, and simulations were run for 1,000 time steps. For a neutral comparison, we also simulated from the same scenario but with no fitness effects. We ran 5 independent simulations of each scenario (i.e., neutral and sexually antagonistic).

After the final generation, genetic load and male-female *F*_*ST*_ at each locus were calculated. *F*_*ST*_ values were calculated both using all individuals within the population as well as using smaller subsamples of 100 individuals and 50 individuals with equal numbers of each sex. Subsample sizes were chosen to reflect sample sizes currently used in the literature (Dutoit *et al.* 2018; Flanagan and Jones 2017). Male-female *F*_*ST*_ values within the subsamples were calculated both with equation 4 and Weir and Cockerham’s *F*_*ST*_ estimator (Weir and Cockerham 1984; Bhatia *et al.* 2013) to examine the impact of the statistic used on the distribution of *F*_*ST*_ values. (With equal female and male subsampling, Weir and Cockerham’s *F*_*ST*_ is equivalent to Hudson’s *F*_*ST*_ (Bhatia *et al.* 2013).)

### Data accessibility

The model equations, equilibrium analyses, and stability analyses are given in the Mathematica notebook in File S1. The single locus simulations and SLiM statistical analyses are given in File S2. The SLiM code and simulation data are available in Files S3-S5. Supplemental material is available at the Genetics Figshare archive: https://gsajournals.figshare.com/s/554868088b2f1eb9a69e.

## Results

We first examine the conditions under which our model supports a stable polymorphism, and then examine the degree of between-sex divergence and genetic load expected under both equilibrium and non-equilibrium (selective sweep) conditions. Our single locus results expand upon previous the results of previous studies (Kidwell *et al.* 1977; Arnqvist 2011; Cheng and Kirkpatrick 2016; Kasimatis *et al.* 2017). We then verify these results using simulations, which also provide an opportunity to explore the effects of statistical sampling on inferences of sex-specific differentiation from genomic samples.

### Transmission dynamics at a sexually antagonistic locus

#### Maintenance of polymorphism usually requires symmetric selection between the sexes under random mating

We will quantify the strength and degree of asymmetry between the sex-specific allelic effects using the overall strength (*s*) and the ratio of selection coefficients (*α*), so that *s*_*m*_ = *s* and *s*_*f*_ = *αs*. The full solution for the maintenance of polymorphism under arbitrary patterns of dominance can be solved by setting *p*(*t*+1) = *p*(*t*) in the recursion equations above (Equations (3); File S1). Under general conditions, this system yields a fifth-order polynomial that does not readily generate a closed form solution in symbolic form, although the equilibria can be easily found numerically. Symbolic solutions are possible under some specific conditions.

Assuming random mating and additivity of allelic effects (*h*_*m*_ = *h*_*f*_ = 0.5), the frequency of the *A*_1_ allele at equilibrium (denoted 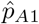) can be expressed in terms of the strength of selection and asymmetry in selection:

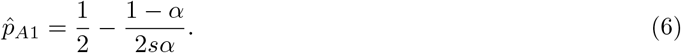

When the strength of selection is equal between the sexes (*α* = 1), an equilibrium frequency of 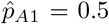 is always predicted. This theoretical solution is well supported by the stochastic simulations as well (Fig. 2A-B). The bounds on the non-trivial equilibrium frequency can be found by setting 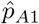 to zero or one. By solving these equations for *α* in terms of the strength of selection (*s*), we find that for the equilibrium to be stable, *α* and *s* must satisfy the condition:

**Figure 2:**
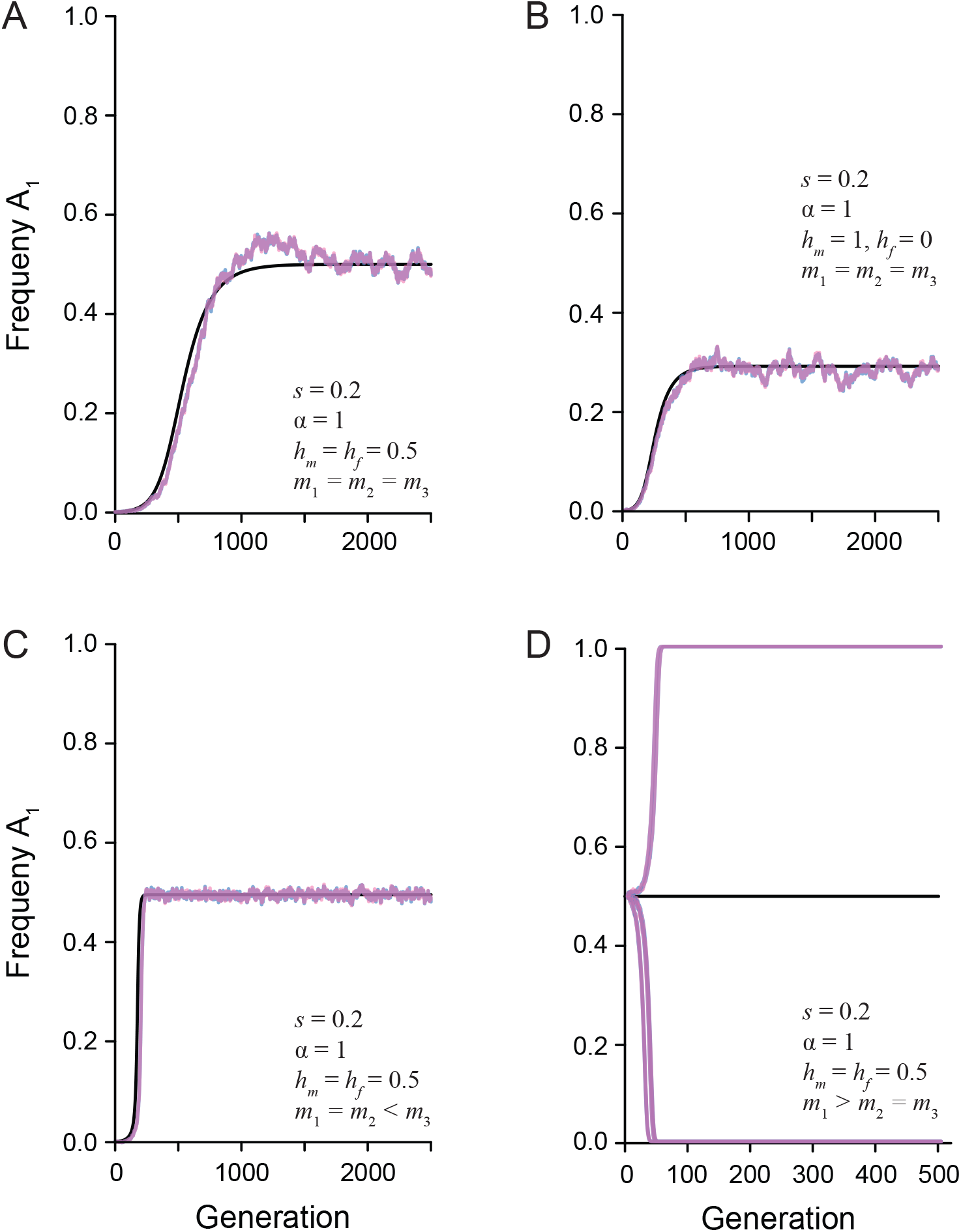
The change in the frequency of a newly derived sexually antagonistic allele (*A*_1_) over time. The black line represents the predicted allele frequency from the recursion equation. The overlaid pink and blue lines represent the simulated population (N = 20,000) of females and males, respectively. The strength of selection (*s*), ratio of selection between the sexes (*α*), dominance relationship (*h*_*m*_ and *h*_*f*_), and mate choice coefficients (*m*_1_, *m*_2_, and *m*_3_) are given in each panel. A) Random mating with additive dominance and symmetric selection between the sexes maintains a stable polymorphism. B) Random mating with complete male dominance and symmetric selection between the sexes maintains a stable polymorphism. C) Assortative mating by fitness with additive dominance maintain a stable polymorphism. D) Assortative mating by genotype with additive dominance has an unstable equilibrium. Multiple simulated populations show how drift will quickly lead to fixation or loss of the *A*_1_ allele.

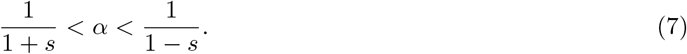

These bounds can also be found by calculating the Jacobian matrix for the full set of transition equations (File S1) and agree with those identified by Kidwell *et al.* (1977). In general, the equilibrium conditions describe an expanding envelope in parameter space that allows more asymmetry in the pattern of antagonistic selection as the absolute strength of selection increases (Fig. 3A & B). To a first order approximation in *s*, equation (7) shows that the equilibrium is stable only if asymmetry is not larger than the strength of selection, such that *|α −* 1*| < s*, as shown in Fig. 3A. Thus, when selection is weak or moderate, the maintenance of a polymorphism requires approximately equal selection between the sexes. However, the permissible degree of asymmetry increases with the strength of selection (Fig. 3B). For example, when *s ≥* 0.4 a stable polymorphism can be maintained so long as the asymmetry in fitness (*|*1 *− α|*) is less than 50%. Selection coefficients of this magnitude mean mortality rates of 40% or higher each generation due to a single incorrect sexually antagonistic allele, which seems biologically implausible. Therefore, under additivity, any stable antagonistic polymorphisms in natural populations must have approximately equal fitness effects in the two sexes, while less balanced antagonistic loci will quickly be fixed or lost.

**Figure 3:**
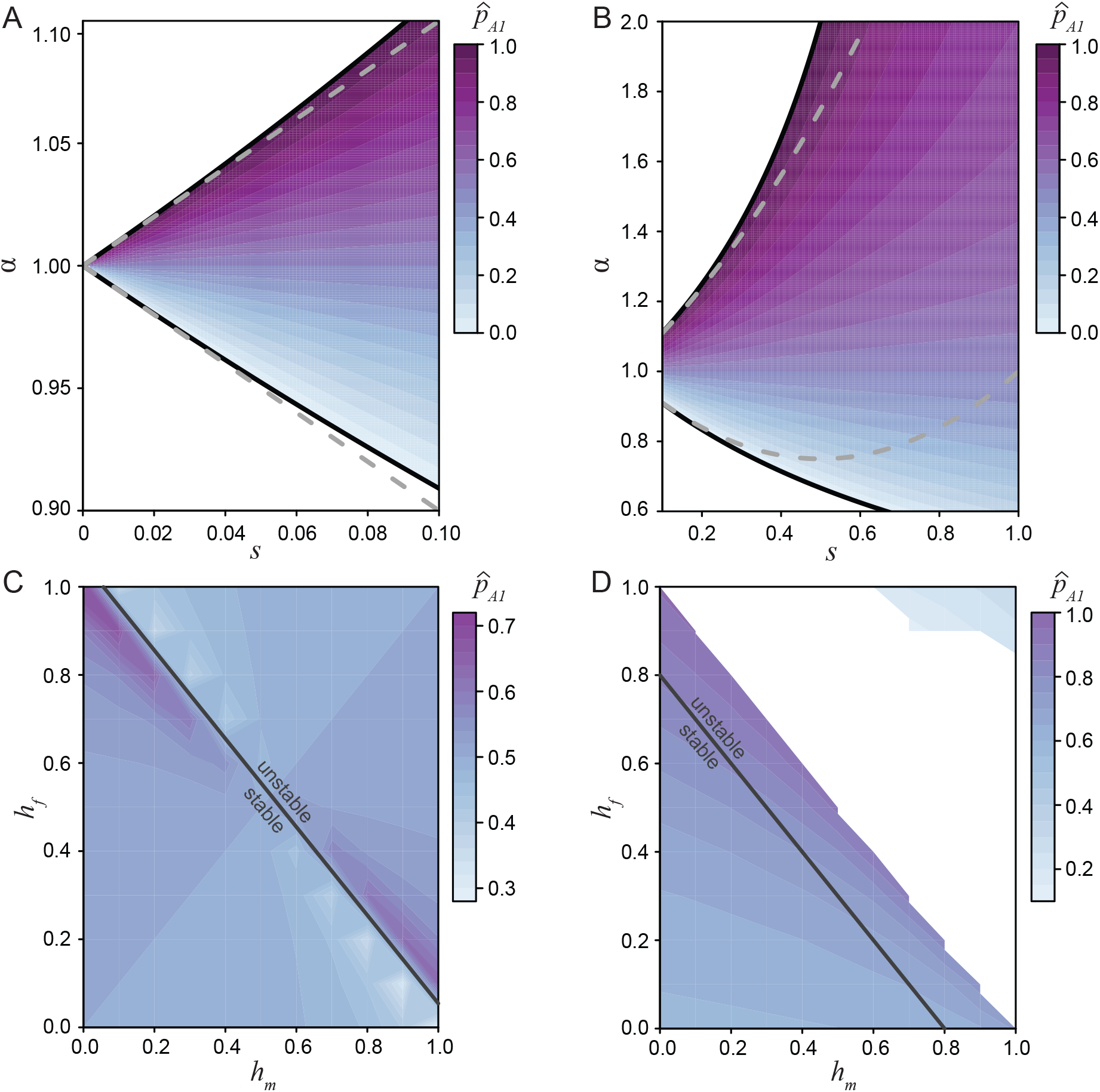
The equilibrium space for the *A*_1_ allele under differing selection and dominance conditions. A) The equilibrium space at an additive locus (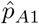, Equation 10), when selection is weak and related between the sexes by the ratio *α*. Here the equilibrium space is symmetric around *α* = 1 and confined to approximately equal selection between the sexes. The solid black line represent the permissible bounds on *α* (7) and the dashed gray line represents the first order Taylor series approximation. B) The equilibrium space at an additive locus increases as the strength of selection increases. The solid black line represents the bounds on *α* and the dashed gray line represents the second order Taylor series approximation. C) The equilibrium space across all dominance conditions when selection is equal between the sexes (*s* = 0.1*, α* = 1). When the dominance coefficients between the sexes sum to no greater than one (*h*_*m*_ + *h_f_ ≤* 1), then the equilibrium is stable. However, when the sum is greater than one the equilibrium is unstable. D) Strong, asymmetric selection (*s* = 0.4*, α* = 1.5) narrows the equilibrium space and range of stable conditions (*h*_*m*_ + *h_f_ ≤* 0.8).

On the other hand, if dominance is allowed to vary between the sexes but selection is equally antagonistic across the sexes (*α* = 1), there is always a single real, non-trivial equilibrium (Fig. 3C), whose stability depends on the sum of the dominance coefficients between the sexes (Kidwell *et al.* 1977). When *h*_*m*_ +*h_f_ ≤* 1 the equilibrium is stable (File S1). This stability boundary makes sense as the mean fitness of homozygous individuals is lower than that of heterozygous individuals (assuming equal sex ratios):

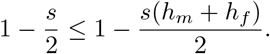

In other words, the equilibrium remains stable if the deleterious effects of dominance in one sex do not outweigh the benefits in the other sex. Interestingly, weak selection at a locus with sex-beneficial dominance (*h*_*m*_ = *h*_*f*_ = 0) can maintain a stable polymorphism despite greater asymmetry in selection than can an additive model (File S1). This expansion of the region of stability is likely a result of heterozygotes being shielded from antagonistic selection and suggests that modifying dominance can act to maintain sexual antagonism at a locus (Connallon and Chenoweth 2019). Conversely, when 1 *< h*_*m*_ + *h_f_ ≤* 2, dominance favors the deleterious allele in each sex, pushing the population to an unstable state and leading to the fixation of the less costly allele. Allowing for asymmetry in the strength of selection narrows the equilibrium space and reduces the range of dominance coefficients resulting in stability (Fig. 3D).

#### Assortative mating by fitness expands the polymorphism space

Under positive assortative mating by fitness, high fitness matings occur between disparate genotypes and therefore produce an excess of heterozygotes each generation. In this situation (i.e., *m*_3_ *> m*_2_ = *m*_1_), up to three real non-trivial equilibria can exist depending on the selection and dominance parameters (File S1). However, as with random mating, at most one equilibrium is stable. When selection is symmetrically antagonistic across the sexes (*α* = 1), an *A*_1_ allele frequency of approximately 0.5 is always predicted, regardless of dominance. This prediction is borne out by the single locus simulation results, which further show that assortative mating by fitness tends to make the stable equilibrium more robust to the effects of genetic drift (Fig. 2C). Increasing the asymmetry of selection can introduce an additional unstable equilibrium, and increasing the strength of sex-deleterious dominance (towards *h*_*m*_ = *h*_*f*_ = 1) can introduce a second unstable equilibrium (File S1). These theoretical predictions agree with previous simulations of assortative mating (Arnqvist 2011). As with random mating, the relationship between the strength and asymmetry in selection is the critical factor in determining when equilibria are stable. Specifically, when the asymmetry in selection is sufficiently large, fixation of the more favored allele is expected. Fixation only tends to occur under unrealistically large viability costs, however, and so the predominant outcome of assortative mating by fitness is the maintenance of heterozygotes and an expansion of the equilibrium space relative to random mating.

#### Assortative mating by genotype leads to fixation

In contrast to assortative mating by fitness, if assortative mating is by genotype (*m*_1_ *> m*_2_ = *m*_3_), there is only a single non-trivial equilibrium (File S1). This equilibrium is always unstable, regardless of dominance, as shown by the leading eigenvalue of the Jacobian matrix. Fig. 2D shows allele frequency trajectories that start at this unstable equilibrium rapidly go to loss or fixation (with the choice determined by random genetic drift). Thus, these mating dynamics shrink the parameter space for maintaining a stable polymorphism and lead to the loss of the weaker antagonistic allele.

### Male-female divergence is exceptionally low

A number of studies have observed high mean male-female divergences (measured by *F*_*ST*_). For instance, Dutoit *et al.* (2018) found a mean male-female *F*_*ST*_ = 0.0016 across genes with male-biased expression in a sample of 43 flycatchers of each sex. Wright *et al.* (2018) found a larger average male-female *F*_*ST*_ value of 0.03 across sex-biased genes in transcriptomes of 11 male and four female Trinidadian guppies. Similarly, Flanagan and Jones (2017) identified 473 genome-wide outliers having male-female *F*_*ST*_ values above roughly 0.05 in a RADseq study of 171 male and 57 female gulf pipefish. Finally, *Lucotte et al.* (2016) found an average male-female *F*_*ST*_ of 0.067 across autosomal SNPs in the human HAPMAP data that showed significant nonzero male-female *F*_*ST*_ in all 11 populations (with around 100 samples of both sexes per population). Previous work showed that selection within a single generation at an additive locus must be strong to generate substantial male-female *F*_*ST*_ values (Cheng and Kirkpatrick 2016; Kasimatis *et al.* 2017). The model we study here allows us to estimate the strength of antagonistic selection required to produce male-female *F*_*ST*_ values as large as these, both at stably polymorphic loci and at loci undergoing a selective sweep.

When selection and dominance coefficients are chosen such that a stable equilibrium is maintained, divergence between the sexes tends to be exceptionally low (Fig. 4A). For example, a 10% viability cost (*s* = 0.1) results in a between-sex *F*_*ST*_ value of 0.0007 at equilibrium (assuming an additive locus and random mating). An equilibrium male-female *F*_*ST*_ value of 0.0016 (as in flycatchers) requires at least a 15% viability cost within each sex (*s* = 0.15, *α* = 1). To produce equilibrium *F*_*ST*_ values an order of magnitude larger (as reported for the largest loci in the other taxa) requires a 30-65% viability cost (*s* = 0.30 *−* 0.65, *α* = 0.8 *−* 2.0). For these values to be a product of viability selection, the field would need to have overlooked as much as 50% genotype-dependant mortality (or infertility) for each sex every generation, which seems implausible in these taxa.

**Figure 4:**
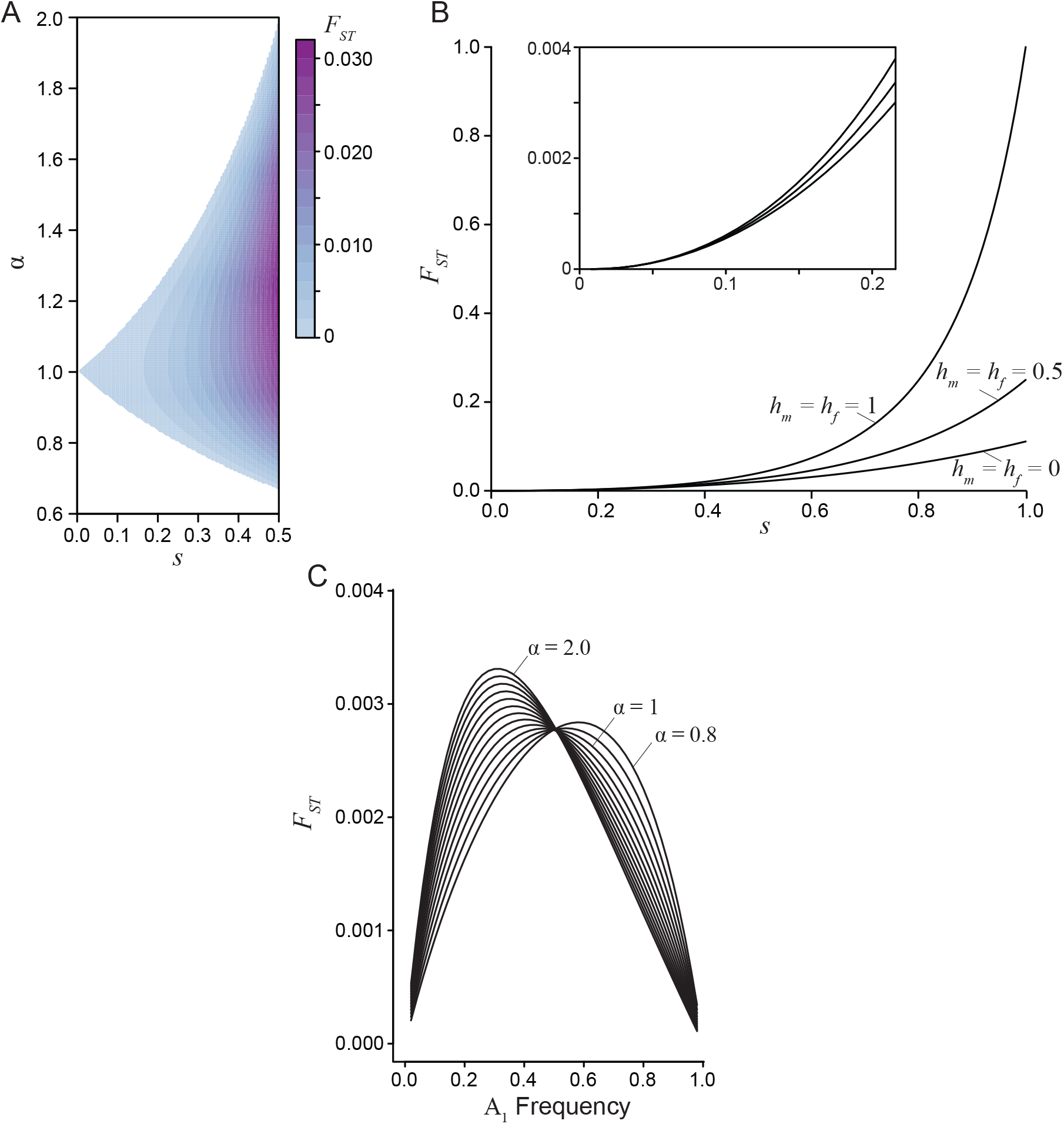
Divergence between the sexes due to a single generation of sexually antagonistic selection. A) Male-female *F*_*ST*_ for an additive locus (*h*_*m*_ = *h*_*f*_ = 0.5) at equilibrium, where the strength of selection between the sexes is related by the ratio *α*. B) Male-female *F*_*ST*_ as a function of selection for three dominance regimes: sex-specific beneficial (*h*_*m*_ = *h*_*f*_ = 0), additive (*h*_*m*_ = *h*_*f*_ = 0.5), and deleterious (*h*_*m*_ = *h*_*f*_ = 1). The sex-specific deleterious dominance curve is at an unstable equilibrium, while the additive and beneficial dominance curves are at a stable equilibrium (for all curves *α* = 1 and the equilibrium frequency is 0.5). Sex-specific beneficial dominance always results in the lowest divergence between the sexes. The inset graph highlights the similarly low divergence values generated under weak and moderately weak selection. C) Male-female *F*_*ST*_ at a sex-beneficial locus (*h*_*m*_ = *h*_*f*_ = 0: the polymorphic equilibrium is stable only if *α* = 1) as a function of *A*_1_ allele frequency for varying degrees of asymmetry in selection (0.8 *≤ α ≤* 2) with a fixed mean selection coefficient (0.5(*s*_*m*_ + *s*_*f*_) = 0.2).

Greater divergence can be generated across a broader range of selection values when an antagonistic locus transiently sweeps through a population. Here a viability cost of 10% produces higher divergence than at equilibrium, although divergence is still low in absolute terms (*F_ST_ <* 0.002 across dominance values, under random mating). Again, at least a 30% viability cost would be required to produce *F*_*ST*_ values above 0.05. Sex-specific beneficial dominance (*h*_*m*_ = 0, *h*_*f*_ = 0) is expected to generate the lowest levels of between-sex divergence, while sex-specific deleterious dominance (*h*_*m*_ = 1, *h*_*f*_ = 1) yields the greatest levels of divergence, though such a scenario seems biologically unstable (Fig. 4B). Importantly, under weak selection dominance has only a negligible effect on divergence. In fact, varying dominance does not generate quantitative changes in *F*_*ST*_ unless selection is remarkably strong (*s >* 0.5). Rather, asymmetry in selection seems a more important driver of divergence in non-equilibrium populations, as this asymmetry is precisely the factor that moves populations away from equilibrium conditions to a state in which the least costly allele sweeps to fixation. Across the range of *α* values with a fixed mean strength of selection between the sexes, divergence slightly increases as asymmetry between the sexes increases (Fig. 4C). However, the male-female *F*_*ST*_ values are of the same magnitude despite strong asymmetry when the mean strength of selection in confined in this manner. When the strength of selection varies independently between the sexes, increasing *α* yields much greater divergence, though this result is confounded by overall stronger selection in one sex. Overall, substantial divergence between the sexes still requires strong selection in non-equilibrium populations.

### Sexual antagonism generates a substantial genetic load

Since the mean relative fitness for females and the mean relative fitness for males are each maximized under fixation for different alleles at an antagonistic locus, sexually antagonistic selection generates a genetic load within the population at both a polymorphic equilibrium and during a selective sweep. At equilibrium under random mating, the load is maximized if the strengths of selection in each sex are equal (Fig. 5A), and dominance has little to no effect. Importantly, across strengths of selection up to *s* = 0.5, the load generated at equilibrium exceeds *F*_*ST*_ between males and females by nearly a factor of 10 (Fig. 5B). For example, a 10% viability cost (*s* = 0.1) results in a reduction of population fitness up to 5%, with a maximum *F*_*ST*_ value of 0.0007. The load produced by a single antagonistic locus with *F*_*ST*_ equal to the **mean** male-female *F*_*ST*_ reported in human HAPMAP data (Lucotte *et al.* 2016) would exceed 20% (Fig. 5B). This relationship indicates that even weak selection driving low – and probably undetectable – levels of divergence can generate a substantial fitness reduction due to the sex-specific nature of selection.

**Figure 5:**
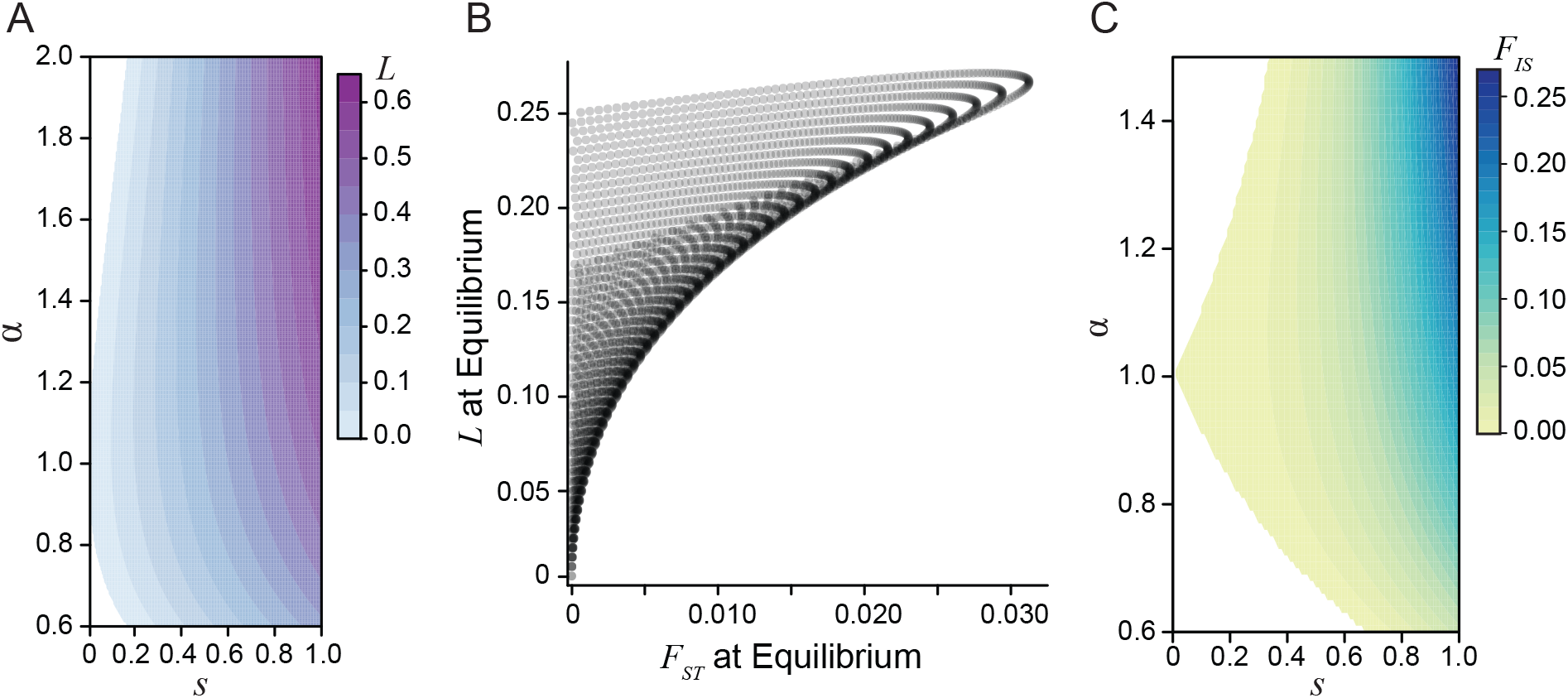
The genetic load created by sexually antagonistic selection. A) The genetic load generated at equilibrium for an additive locus across strengths (*s*) and asymmetries (*α*) of selection. B) A comparison of male-female divergence and genetic load for a locus at equilibrium across a gradient of selection coefficients with varying asymmetry. The load generated at a locus exceeds the degree of divergence between the sexes. Each curve corresponds to a different fixed strength of selection from *s* = 0 to *s* = 0.5 and each point along the curves corresponds to a different value of *α* from 0.6 to 2. C) The population inbreeding coefficient *F*_*IS*_ for an additive locus (*h*_*m*_ = *h*_*f*_ = 0.5) at equilibrium. The excess of heterozygous individuals in the population represents the departures from Hardy-Weinberg equilibrium due to sex-specific selection.

An alternative way to examine load is by quantifying the excess of heterozygosity due to sexually antagonistic selection, using the *F*_*IS*_ statistic (the inbreeding coefficient). Here sex-specific selection creates homozygous pools of each sex within a generation, which leads to an excess of heterozygotes at the start of the next generation. Under weak selection, these departures from Hardy-Weinberg equilibrium are of similar magnitude as male-female *F*_*ST*_ (Fig. 5C). However, under strong sexual antagonism – such as that required to generate the empirically observed divergence values – *F*_*IS*_ can approach 10%.

Antagonistic loci that do not have a polymorphic equilibrium tend to produce even greater load while sweeping than a locus at stable equilibrium under similar selective conditions (File S1). Here load was approximately half of the strength of selection for the allele with the higher fitness cost (i.e., *sα/*2). Under strong, asymmetric selection load can approach 50% during a sweep. Additionally, the fitness cost of sexual antagonism remains after an allele fixes. The load generated during a sweep is affected by dominance, with additive loci generating loads that are intermediate to the other dominance scenarios. Beneficial dominance within each sex can apparently resolve some of the underlying antagonism by shielding selection on heterozygotes and therefore reducing the load. In contrast, sex-specific deleterious dominance generated the greatest load.

### Genome-wide antagonistic selection also produces low divergence

Our analytical results are based on a single-locus model, yet empirical studies report averages across large numbers of loci. To complement the single-locus theory, we quantified the effects of sexually antagonistic selection throughout the genome using individual-based simulations in SLiM Haller and Messer (2019). First, we analyzed male-female *F*_*ST*_ values calculated using true population-wide allele frequencies (i.e., whole-population *F*_*ST*_). Simulations in which every new mutation was sexually antagonistic in a population of 10,000 individuals resulted in a mean male-female *F*_*ST*_ of 0.00005 and a between-replicate standard deviation of 0.0001, consistent with the single-locus theory (since *s* was around 0.01). However, entirely neutral simulations (equal mutation rates but no selection) resulted in the same mean and SD of male-female *F*_*ST*_ values. Both the sexually antagonistic and neutral simulations averaged around 1,400 SNPs after 1,000 generations of evolution. Although qualitatively similar, the distribution of whole-population male-female *F*_*ST*_ values across loci was statistically significantly different between the neutral and sexually antagonistic simulations (Kolmogorov-Smirnov test: *D* = 0.11, *p <* 0.001; Fig. 6A). However, this difference in distributions was driven by the larger number of intermediate frequency alleles in the sexually antagonistic simulations. In particular, neutral simulations across all five replicates had only two SNPs with a frequency above 10%, while the sexually antagonistic simulations had over 200 SNPs with a frequency above 10%. Despite there being true differences between the neutral and sexually antagonistic simulations, the male-female divergences observed were still exceptionally low. In fact, neither model had any loci with male-female *F*_*ST*_ greater than 0.001 (Fig. 6A). Sexually antagonistic simulations had an average 21% decrease in population fitness (*L* = 0.21 *±* 0.02) after 1000 generations of evolution, again consistent with single-locus calculations. Even the minimum load observed under the sexually antagonistic scenario corresponded to a 18% decrease in population fitness.

**Figure 6:**
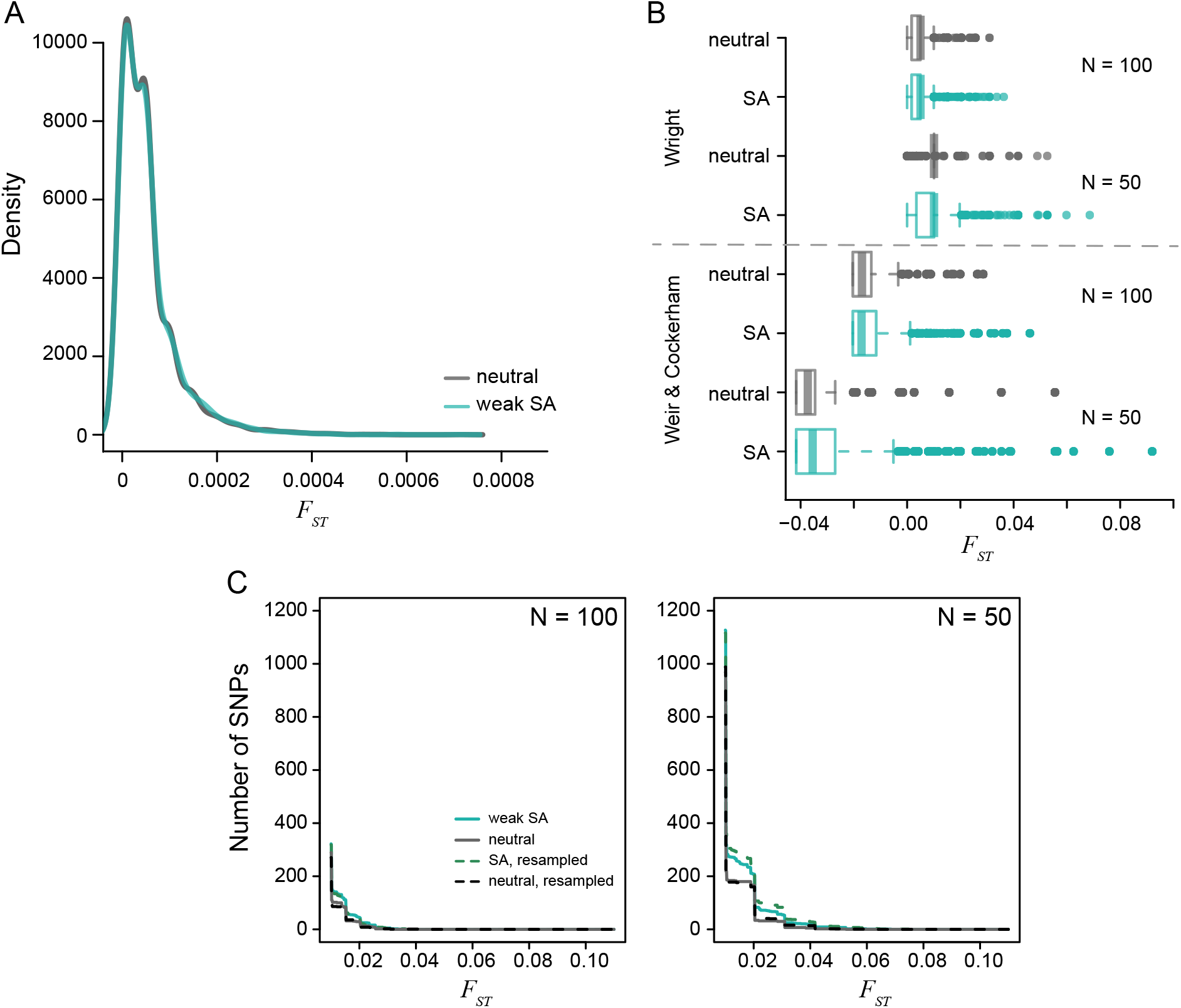
The distribution of per locus *F*_*ST*_ values generated from simulated populations after 1000 generations of evolution. A) The density of male-female *F*_*ST*_ values for a population of 10,000 individuals is centered around *F*_*ST*_ = 0.0005. The neutral (gray) and sexually antagonistic (teal) simulations were similar but statistically significantly different. B) The distribution of male-female *F*_*ST*_ values when subsampling the full populations to either 100 individuals or 50 individuals with equal sex ratios. Divergence values were calculated using the theoretical derivation (Eq. 4) and Weir and Cockerham’s *F*_*ST*_. Subsampling increases the tail of the distribution substantially. The sexually antagonistic simulations are significantly different from the neutral simulations due to an increased sampling variance in the sexually antagonistic scenario. C) Cumulative distribution curves of per-locus male-female *F*_*ST*_ values, both between random samples from the two sexes (solid lines) and between sets of individuals chosen randomly independently of sex (dotted lines). Male-female *F*_*ST*_ distributions differed between neutral (grey) and antagonistic (teal) simulations but were not higher for between-sex comparisons, showing that higher *F*_*ST*_ values in the antagonistic simulation was not directly due to selection.

### Sampling variance can generate spurious signals of male-female divergence

Both the single locus model and genome-wide simulations indicate that, while theoretically possible, we would need strong sexually antagonistic viability selection to maintain high divergence between the sexes. Alternatively, the large observed *F*_*ST*_ statistics might be due to sampling variance. The empirical studies we cite have relatively small sample sizes (between 15 and 200). The male-female *F*_*ST*_ values we reported above from simulation were calculated from the *entire* population. To evaluate the effect of sampling, we calculated male-female *F*_*ST*_ values from random samples of individuals in our SLiM simulations of two sizes: 100 individuals (50 females and 50 males) and 50 individuals (25 females and 25 males). This subsampling produced dramatically higher male-female *F*_*ST*_ values under both the neutral (100 individuals, mean *±* standard deviation across replicates: *F*_*ST*_ = 0.005 *±* 0.004; 50 individuals: *F*_*ST*_ = 0.01 *±* 0.007) and sexually antagonistic (100 individuals: *F*_*ST*_ = 0.005 *±* 0.004; 50 individuals: *F*_*ST*_ = 0.01 *±* 0.008) simulations (Fig. 6B). Additionally, the tail of the distribution was lengthened under both neutral and sexually antagonistic simulation, including in both cases estimated values of *F*_*ST*_ several orders of magnitude larger than the values calculated from actual population allele frequencies. In other words, ignoring the noisiness of the *F*_*ST*_ estimator would lead us to conclude that between-sex divergence is much higher than it actually is. There was a significant difference in the distribution of male-female *F*_*ST*_ values between the neutral and sexually antagonistic simulations both when subsampling at 100 individuals (Kolmogorov-Smirnov test: *D* = 0.054, *p <* 0.01) and 50 individuals (*D* = 0.096, *p <* 0.001). However, there was no correlation between the *F*_*ST*_ values calculated from the full population and those obtained from samples of either 100 individuals (*r* = 0.003) or the 50 individual subset (*r* = *−*0.012). This lack of correlation also holds true for the neutral model (100 individuals: *r* = 0.034; 50 individuals: *r* = 0.025).

Empirical studies often use estimators of *F*_*ST*_, such as Weir and Cockerham’s *F*_*ST*_ to account for population size and sample allele frequencies, rather than population allele frequencies. Therefore, in addition to using equation 4, we calculated male-female *F*_*ST*_ values in subsamples using Weir and Cockerham’s method (Weir and Cockerham 1984; Bhatia *et al.* 2013). However, this did not qualitatively affect the tail of the distribution of *F*_*ST*_ (Fig. 6B). This is not surprising, because Weir and Cockerham’s method is designed to obtain *unbiased* estimates of *F*_*ST*_ – and so re-centers the mean of the distribution – but does not reduce sampling variance.

Although there were more high male-female *F*_*ST*_ sites in samples from the sexually antagonistic simulations (Fig. 6B), this did not seem to be a direct result of selection, but was due to the fact that there are many more intermediate frequency alleles in the sexually antagonistic simulations because of balanced polymorphisms. (If the larger number of high *F*_*ST*_ sites were due to selection on those sites, then we’d expect to see a correlation between estimated *F*_*ST*_ and population *F*_*ST*_, which, as noted above, we do not.) To test this hypothesis, we calculated *F*_*ST*_ between two random samples of size 50 drawn from each simulation *independently* of sex, and also between random samples of size 25. If the enrichment of high between-sex *F*_*ST*_ sites in the antagonistic simulations are in fact due to the difference in allele frequency distribution rather than the direct result of selection, then the enrichment should persist even in these samples drawn after randomizing sex. Indeed this enrichment persists, as shown in Fig. 6C. Thus, the higher number of intermediate frequency sites in the sexually antagonistic model creates a higher sampling variance of *F*_*ST*_, as expected based on theory (Jakobsson *et al.* 2013). In particular, the tail of the *F*_*ST*_ distributions show many higher values, such that an increase by two orders of magnitude relative to the full population was observed (Fig. 6B). These results suggest that separating signals of weak antagonistic selection from sampling noise will be extremely difficult.

## Discussion

Sexually antagonistic viability selection creates allelic divergence between the sexes because the proportions of each genotype that die before reproduction differs between the sexes. This between-sex divergence for non-sex-linked elements is created anew each generation because chromosomal segregation re-assorts autosomal associations across the sexes during sexual reproduction. An emerging trend in sexual antagonism research is the use of male-female genomic comparisons to identify sexually antagonistic loci. These recent studies identified hundreds of sexually divergent autosomal loci with mean divergence between the sexes in the range of 2-7% (Lucotte *et al.* 2016; Flanagan and Jones 2017; Wright *et al.* 2018). Taken as reported, these studies suggest the extent and strength of sexually antagonistic selection is far greater than might be anticipated. To assess these claims, we used a population genetic model to determine the magnitude of divergence generated by sexually antagonistic viability selection, the strength of selection required to drive such divergence, and the population fitness costs generated by this process.

Although sexual antagonism has been a topic of particular interest over the last few decades (Arnqvist and Rowe 2005), some of the early investigations of sex-specific selection were largely motivated as part of a general attempt to elucidate all possible means by which the large amounts of segregating polymorphisms observed within natural populations could be maintained (Lewontin 1974). In this context, *Kidwell et al.* (1977) focused on how sex-specific selection, could maintain a polymorphism at an autosomal locus when alleles had opposing effects in the sexes. Our analysis agrees with Kidwell *et al.* (1977), though we highlight that maintenance of such polymorphisms would create a substantial genetic load. Additionally, with weaker selection, the parameter space allowing a stable polymorphism becomes quite narrow. These results highlight the necessity for considering biologically relevant conditions – as discussed by Smith and Hoekstra (1980) – particularly when theory is is informing signatures of selection within the genome. Our analysis also allows for non-random mate choice, a potentially important underlying component of sexual conflict (Arnqvist and Rowe 2005). We build on Bodmer (1965) to generate a fully generalizable mate choice matrix and find that assortative mating can indeed have a large impact on the conditions for the maintenance of polymorphism. Supporting previous simulations (Arnqvist 2011), we found positive assortative mating by fitness maintained polymorphisms. In particular, we show that the combination of asymmetrical selection between the sexes and deleterious dominance conditions expanded the equilibrium space relative to random mating. However, such deleterious sex-specific dominance would likely be selected against, suggesting that the strength of selection is the more relevant parameter in natural populations.

While the maintenance of polymorphism may have been a primary motivation for previous work, a goal of modern genomics is to use specific signals of genomic differentiation to identify the loci underlying sexually antagonistic genetic effects (Mank 2017a). Building on our previous work (Kasimatis *et al.* 2017) allowed us to consider the expected degree of between-sex divergence both when an antagonistic polymorphism is maintained at equilibrium in the population, and when no such stable equilibrium exists, so one of the two alleles sweeps towards fixation to the detriment of one sex. Our model and accompanying simulations highlight several potential limitations of detecting sex-specific differentiation in empirical studies.

First, detectable quantitative divergence between the sexes requires exceptionally strong sexually antagonistic selection. Previous work indicates that *F*_*ST*_ values between populations is of order *s*^2^ (Charlesworth and Charlesworth 2010; Cheng and Kirkpatrick 2016), but we find that substantially lower values of *F*_*ST*_ are often obtained in practice. Even a 10% viability cost in each sex resulted in between-sex *F*_*ST*_ values of less than 0.001 (Fig. 4), a signal that is unlikely to be distinguishable from noise without sampling many thousands of individuals within each sex. Critically, to achieve divergence values greater than 0.03 – such as seen in estimates from human data – would require a 30% to 60% viability cost in each sex under our model. These remarkably high sex-specific mortality rates are, to the best of our knowledge, not observed in nature (see Singh and Punzalan 2018). (However, exceptionally high fecundity animals might withstand such high sex-specific mortality (see Williams 1975).) Cheng and Kirkpatrick (2016) also pointed out that even small male-female *F*_*ST*_ values would require unrealistic amounts of genetic load, but still argue in favor of ongoing sex-specific selection at many genes. Strong sex-specific gametic selection is perhaps more plausible than viability selection on adults, but such high levels of genotype-dependent gamete “mortality” in these organisms still do not seem consistent with empirical observations. Additionally, gametic selection requires an epistatic association between the autosomal locus and the sex determining region, which seems highly implausible across so many loci.

Second, asymmetry in the strength of selection between the sexes is critical in determining the degree of divergence generated. When the strength of selection is weak and approximately the same between the sexes, polymorphisms may be stably maintained, but between-sex divergence is small. However, there is no *a priori* reason to expect that antagonistic mutations should be perfectly symmetrical in their effects and therefore that polymorphic loci should be stable over time. Alleles with more asymmetric effects will often sweep, producing larger but transient between-sex divergences, although again only under moderate to strong selection. Of course, other types of selection (e.g., spatially varying selection) could contribute to maintenance of more asymmetric polymorphisms, but the observations about genetic load should still hold. Here, we found that dominance has little quantitative effect on male-female divergence, particularly when selection is weak. In general, understanding what the distribution of sex-specific effects underlying antagonistic selection looks like will provide important information on the potential for sexually antagonistic loci to contribute to genetic variation and genome evolution.

Thus, the theoretical predictions from our single locus model seem at odds with the empirical patterns reported to date. Taken as true measurements of sexually antagonistic selection, the empirical data could be described by two, non-exclusive genomic patterns. Divergent loci could either be stable polymorphisms or could be arising and sweeping to fixation through a constant genomic churn of antagonistic interactions. Either of these explanations require an exceptionally high genetic load. Again, there is currently no indication that mortality occurs in such a high, sex-specific manner, particularly in some of the vertebrate species that have been examined.

Individual-based simulations with many linked selected loci genome-wide recapitulate the predictions of the single-locus model, finding again that even in this more complex situation, weak selection can only produce very low levels of divergence. Most importantly, however, we found that estimating male-female *F*_*ST*_ from samples of the sizes used in the literature (hundreds or less) produced distributions with larger means and longer tails, even in the complete absence of antagonistic selection. Even in simulations with antagonistic selection, any high divergence values were a result of random sampling noise, and did not correlate with the true divergence values or strength of selection. These simulations highlight the sensitivity of *F*_*ST*_ statistics to sampling variance, which is a major obstacle for identification of antagonistic loci from sex-specific differentiation. Most existing empirical studies have not taken these effects fully into account. Our simulations are not intended to be comprehensive, but demonstrate that sampling variance can be more important than selection itself in driving high estimates of divergence, and highlight the need for proper sampling theory.

At the very least, studies analyzing male-female *F*_*ST*_ values should compare values to empirical distributions found by random permutation of sex labels, as done by Dutoit *et al.* (2018). Connecting significant SNPs to a phenotype such as sex-biased expression (Cheng and Kirkpatrick 2016; Dutoit *et al.* 2018) can provide additional evidence for selection, but it is difficult to control for all confounding factors (e.g., overall expression level). Furthermore, population substructure remains a concern even when comparing to permuted data if there is a cryptic correlation of sampling with sex. For instance, suppose that the sampled population is composed of a mixture of two diverged subpopulations, and that the sex and admixture coefficients of the sampled individuals are correlated. As noted by Cheng and Kirkpatrick (2016), this will create spurious male-female *F*_*ST*_. (The samples in Dutoit *et al.* (2018) were composed of mating pairs from a single island, so this seems unlikely to explain their results, unless one sex is much more likely to disperse between islands than the other.) Other issues beyond sampling variance may well play a role in the large observed male-female *F*_*ST*_ values. Reads from the sex chromosome that are wrongly aligned to an autosome, particularly in the heterogametic sex, have the potential to generate spurious *F*_*ST*_ peaks (see *Tsai et al.* (2019)), an issue that may affect some classes of genes – such as those with sex-biased expression – more than others.

Furthermore, some studies report large numbers of loci with high average *F*_*ST*_. Should we interpret this as evidence of antagonistic selection across many loci simultaneously, or at just a few loci that affect others through linkage? This is not clear, because each generation’s sex-specific selection on a single antagonistic allele will also cause between-sex frequency differences at other loci to the extent they are in linkage disequilibrium with the locus under selection. More work is needed to quantify this effect.

More generally, our investigation calls into question whether it is even possible, at any sample size, to identify the action of realistically strong sex-specific antagonistic selection using male-female *F*_*ST*_. Many of the loci previously suggested as contributing to sex-specific antagonistic selection seem likely to be spurious signals resulting from poor statistical inference. While we believe sexually antagonistic selection does contribute to genomic evolution, we strongly caution against the use and over-interpretation of male-female *F*_*ST*_ statistics, especially with small sample sizes, and until the potential bioinformatic confounders are better understood.

## Acknowledgements

We thank Göran Arnqvist and Thomas Nelson for their helpful discussion and two anonymous reviewers for their comments. This work was supported by the National Institutes of Health (training grant T32GM007413 to KRK and R01GM102511 to PCP) and the ARCS Oregon Chapter (KRK).

## References

Arnqvist, G., 2011 Assortative mating by fitness and sexually antagonistic genetic variation. Evolution 65: 2111–2116.

Arnqvist, G. and L. Rowe, 2005 Sexual conflict. Princeton University Press, Princeton, NJ.

Bhatia, G., N. Patterson, S. Sankararaman, and A. L. Price, 2013 Estimating and interpreting FST: the impact of rare variants. Genome Research 23: 1514–1521.

Bodmer, W. F., 1965 Differential fertility in population genetics models. Genetics 51: 411–424.

Bonduriansky, R. and S. F. Chenoweth, 2009 Intralocus sexual conflict. Trends Ecol. Evol. 24: 1–9.

Charlesworth, B. and D. Charlesworth, 2010 Elements of evolutionary genetics. Roberts & Company, En-gelwood, CO.

Charlesworth, D. and B. Charlesworth, 1980 Sex differences in fitness and selection for centric fusions between sex-chromosomes and autosomes. Genet. Res. 35: 205–214.

Cheng, C. and M. Kirkpatrick, 2016 Sex-specific selection and sex-biased gene expression in humans and flies. PLoS Genet. 12: e1006170–18.

Connallon, T. and S. F. Chenoweth, 2019 Dominance reversals and the maintenance of genetic variation for fitness. PLoS Biol. 17: e3000118.

Connallon, T. and A. G. Clark, 2011 The resolution of sexual antagonism by gene duplication. Genetics 187: 919–937.

Connallon, T., R. M. Cox,and R. Calsbeek, 2010 Fitness consequences of sex-specific selection. Evolution 64: 1671–1682.

Dutoit, L., C. F. Mugal, P. Bolívar, M. Wang, K. Nadachowska-Brzyska, et al., 2018 Sex-biased gene expression, sexual antagonism and levels of genetic diversity in the collared flycatcher (ficedula albicollis) genome. Mol. Ecol. pp. 1–31.

Flanagan, S. P. and A. G. Jones, 2017 Genome-wide selection components analysis in a fish with male pregnancy. Evolution 71: 1096–1105.

Fry, J. D., 2010 The genomic location of sexually antagonistic variation: some cautionary comments. Evolution 64: 1510–1516.

Haldane, J. B. S., 1937 The effect of variation of fitness. Am. Nat. 71: 337–349.

Haller, B. C. and P. W. Messer, 2019 SLiM 3: forward genetic simulations beyond the Wright-Fisher model. Mol. Biol. Evol. 36: 632–637.

Immler, S. and S. P. Otto, 2018 The evolutionary consequences of selection at the haploid gametic stage. Am. Nat. 192: 241–249.

Jaenike, J., 2003 Sex chromosome meiotic drive. Ann. Rev. Ecol. Systemat. 32: 25–49.

Jakobsson, M., M. D. Edge, and N. A. Rosenberg, 2013 The relationship between FST and the frequency of the most frequent allele. Genetics 193: 515–528.

Joseph, S. and M. Kirkpatrick, 2004 Haploid selection in animals. Trends Ecol. Evol. 19: 592–597.

Kasimatis, K. R., T. C. Nelson, and P. C. Phillips, 2017 Genomic signatures of sexual conflict. J. Hered. 108: 780–790.

Kidwell, J. F., M. T. Clegg, F. M. Stewart,and T. Prout, 1977 Regions of stable equilibria for models of differential selection in the two sexes under random mating. Genetics 85: 171–183.

Kozielska, M., F. J. Weissing, L. W. Beukeboom,and I. Pen, 2010 Segregation distortion and the evolution of sex-determining mechanisms. Heredity 104: 100–112.

Lewontin, R. C., 1974 The genetic basis of evolutionary change. Columbia University Press, New York.

Lucotte, E. A., R. Laurent, E. Heyer, L. Ségurel,and B. Toupance, 2016 Detection of allelic frequency differences between the sexes in humans: a signature of sexually antagonistic selection. Genome Biol. Evol. 8: 1489–1500.

Mank, J. E., 2017a Population genetics of sexual conflict in the genomic era. Nature Rev. Genet. 7: 1–10.

Mank, J. E., 2017b The transcriptional architecture of phenotypic dimorphism. Nature Ecol. Evol. 1: 1–7.

Patten, M. M. and D. Haig, 2009 Maintenance or loss of genetic variation under sexual and parental antagonism at a sex-linked locus. Evolution 63: 2888–2895.

R Core Team, 2018 R: A Language and Environment for Statistical Computing. R Foundation for Statistical Computing, Vienna, Austria.

Rice, W. R., 1984 Sex chromosomes and the evolution of sexual dimorphism. Evolution 38: 735–742.

Rice, W. R., 1987 The accumulation of sexually antagonistic genes as a selective agent promoting the evolution of reduced recombination between primitive sex chromosomes. Evolution 41: 911–914.

Rice, W. R. and B. Holland, 1997 The enemies within: intergenomic conflict, interlocus contest evolution (ICE), and the intraspecific Red Queen. Behav. Ecol. Sociobiol. 41.

Singh, A. and D. Punzalan, 2018 The strength of sex-specific selection in the wild. Evolution.

Smith, J. M. and R. Hoekstra, 1980 Polymorphism in a varied environment: how robust are the models? Genet. Res. 35: 45–57.

Tsai, K. L., J. M. Evans, R. E. Noorai, A. N. Starr-Moss, and L. A. Clark, 2019 Novel Y Chromosome Retrocopies in Canids Revealed through a Genome-Wide Association Study for Sex. Genes 10: 320–11.

Weir, B. S. and C. C. Cockerham, 1984 Estimating F-statistics for the analysis of population structure. Evolution 38: 1358–1370.

Williams, G. C., 1975 Sex and evolution.. Princeton University Press, Princeton, NJ.

Wright, A. E., M. Fumagalli, C. R. Cooney, N. I. Bloch, F. G. Vieira, et al., 2018 Male-biased gene expression resolves sexual conflict through the evolution of sex-specific genetic architecture. Evol. Letters 215: 403–10.

Wright, S., 1951 The genetical structure of populations. Annals of eugenics 15: 323–354.

